# Early life DNA methylation profiles are indicative of age-related transcriptome changes

**DOI:** 10.1101/557892

**Authors:** Niran Hadad, Dustin R. Masser, Laura Blanco-Berdugo, David R. Stanford, Willard M. Freeman

## Abstract

Alterations to cellular and molecular programs with brain aging result in cognitive impairment and susceptibility to neurodegenerative disease. Changes in DNA methylation patterns, an epigenetic modification required for various CNS functions, are observed with aging and can be prevented by anti-aging interventions, but the functional outcomes of altered methylation on transcriptome profiles are poorly understood with brain aging. Integrated analysis of the hippocampal methylome and transcriptome with aging of male and female mice demonstrates that age-related differences in methylation and gene expression are anti-correlated within gene bodies and enhancers, but not promoters. Methylation levels at young age of genes altered with aging are positively associated with age-related expression changes even in the absence of significant changes to methylation with aging, a finding also observed in mouse Alzheimer’s models. DNA methylation patterns established in youth, in combination with other epigenetic marks, are able to predict changes in transcript trajectories with aging. These findings are consistent with the developmental origins of disease hypothesis and indicate that epigenetic variability in early life may explain differences in age-related disease.

## Introduction

Epigenetic modifications, chromatin and direct DNA modifications, are key genomic regulatory processes required for proper development [1], gene imprinting [2-4], X chromosome inactivation [5-7], gene expression regulation [8], and genomic organization [9-11]. Disruptions to the epigenome can alter basic cellular regulation leading to a wide range of dysfunctional molecular programs [10-12]. Impaired epigenetic control with aging has been proposed as an etiological factor common to age-related diseases ranging from diabetes to neurodegenerative diseases such as Alzheimer’s disease [13-18]. DNA methylation has been widely studied in geroscience research as methylation at specific loci tracks with chronological aging [19-22] and can potentially be an indicator of ‘biological’ aging [23,24]. DNA methylation primarily occurs in a CpG context, however non-CpG methylation is abundant in the central nervous system (CNS) [1,25] and has only been minimally examined with aging[26,27]. With the growing understanding that DNA methylation is dynamic, the role of alterations in DNA methylation patterns in regulating gene expression changes during development, aging, and disease is of particular interest.

DNA methylation changes with aging demonstrate both tissue specificity and conservation across tissues depending on the specific genomic location [28-30], with hypermethylation of the ELOVL2 promoter being one of the most well described [31,32]. Conserved changes with aging in the form of epigenetic clocks have proved to be a powerful tool for estimating chronological age and are predictive of all-cause mortality [24,33,34]. Tissue specific DNA methylation changes with aging on the other hand may underlie organ/cell specific deficits. For example, in the liver, gene body hypermethylation occurs primarily in genes involved in lipid metabolism [35], while in the brain age-related methylation changes occur in genes involved in synaptic transmission and cellular integrity [26]. It is important to note that changes in methylation also occur in pathways implicated to be dysregulated with aging systemically, such as the insulin signaling pathway and cellular senescence [36-39]. Recent studies show that age-related DNA methylation changes in blood [40,41], kidney [42], liver [35,39], and the hippocampus [26], can be prevented by dietary, genetic, and pharmacological pro-longevity interventions providing further support for the association between DNA methylation and aging.

In the CNS, DNA methylation plays an important role in cellular differentiation [43-45], synaptic formation and function [46,47], and in molecular mechanisms underlying learning and memory formation [48]. These processes are known to be impaired with aging [49]; however, whether age-related methylation differences contribute to the decline of these processes is unknown. Global levels of DNA methylation have been proposed to decrease with aging [50], but have not been found to be altered with age in the brain using modern sequencing techniques [51,52]. Rather specific loci in the genome undergo hypermethylation and hypomethylation with aging [27]. In addition to differences in methylation, with aging there is increased variability in CpG methylation [53]. Similar findings are observed in Alzheimer’s disease (AD) patients, specifically in genes directly linked to AD [17]. These observations contribute to the idea that epigenetic mechanisms may contribute to age-related impairments and disease through altering gene expression. Yet, little is known about the effects of age-related changes in methylation on gene expression regulation in the brain. Understanding the role age-related differential methylation plays in brain aging may allow for identification of regulatory processes contributing to the development of neuropathologies.

In previous studies we have characterized changes in methylation and transcription with aging in the hippocampus of male and female mice, finding a core of sex common changes with the majority of age-related changes being sexually divergent [27,54]. Here we sought to understand the effect of age-related differential methylation on gene expression using paired DNA methylation, by whole-genome bisulfite sequencing (WGBS), and transcriptome, by RNA-sequencing, data from the same samples. We find that differential methylation in gene body and enhancer elements inversely correlates with aging gene expression. This relationship is generally weak and accounts for a small fraction of the differentially expressed genes with aging. A stronger correlation was observed between age-related differential gene expression and methylation patterns that are already established in young animals, an association that was independent of age-related differential methylation. Furthermore, DNA methylation levels were able to predict whether transcriptional changes with age will undergo up- or down-regulation with aging. The predictive ability increased when combined with other epigenetic marks. The broad implication of our findings suggests that early programming of the epigenome during development and/or early adulthood may impact transcriptional trajectories late in life. Understanding epigenetic differences that occur during development may help explain late-life molecular responses in the CNS and possibly differences in susceptibility to adverse conditions between individuals.

## Results

### Characterization of differential methylation in the hippocampus using whole genome bisulfite sequencing

To assess the relationship between hippocampal age-related differential methylation and age-related transcriptional changes we first analyzed differential methylation with aging using WGBS. Previous studies characterizing differential methylation in the hippocampus with aging focused on global levels of methylation or used approaches that allowed for high resolution analysis of portions (∼10%) of the genome [27,51]. Whole genome bisulfite sequencing provides a complete analysis of gene methylation by covering the majority of CpG sites across the genome. Sequencing methods that examine smaller portions of the CpG sites in the genome provide limited and incomplete information regarding genic methylation (Supplemental Figure 1).

**Figure 1:**
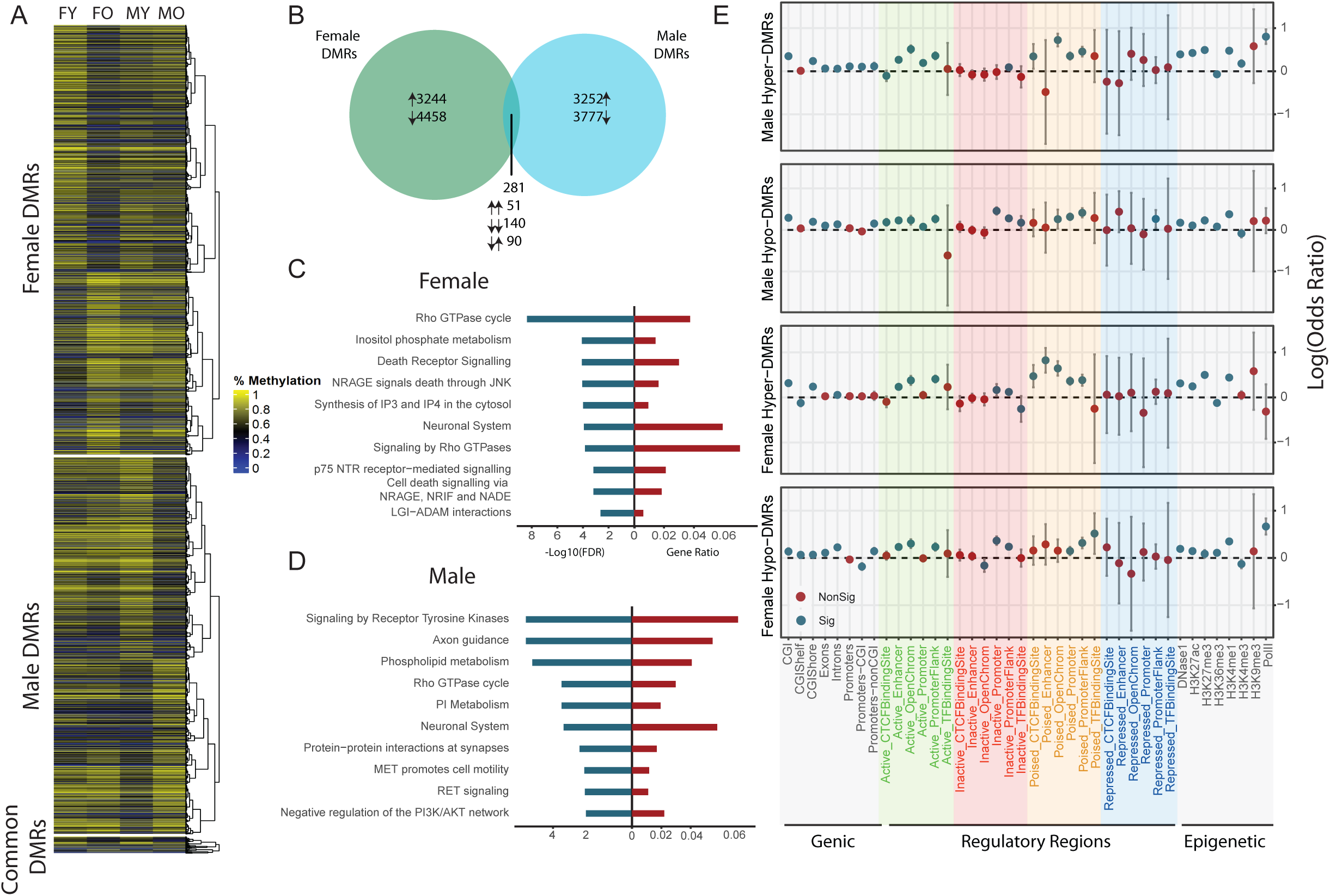
Whole genome characterization of age-related differential methylation with aging in males and females. A) Heatmap showing methylation values of age-related differentially methylated regions (Fisher Exact Test with FDR < 0.05, n=3/group) across all groups. B) Venn diagram showing overlap between age-DMRs in males and in females and the directionality of methylation changes of common age-DMRs. C,D) Pathway enrichment of genes containing age-DMR within their gene body in females (C) and in males (D). Significant enrichment was determined by hypergeometric test (p<0.05). E) Over- and under-representation of age-DMRs in genic regions, CpG islands, regulatory elements in the brain divided by their activation state, and regulatory elements annotated by specific histone marks in males and females, separated by hyper- and hypo-methylation. Over- and under-representation was determined using hypergeometric test (p< 0.05).

The average methylation level across all CpGs in young (3 months) and old (24 months) animals demonstrate no difference with aging (FY 74% ±0.2, FO 73.5% ±0.4, MY 74.1% ±0.5, MO 72.5% ±1.4, Supplemental Figure 2). Similarly, no difference in transposable element CpG methylation with age was evident. Differences in average methylation levels were also not observed between males and females. These agree with previous findings that there is no hypomethylation with aging in the murine hippocampus [51,52].

**Figure 2:**
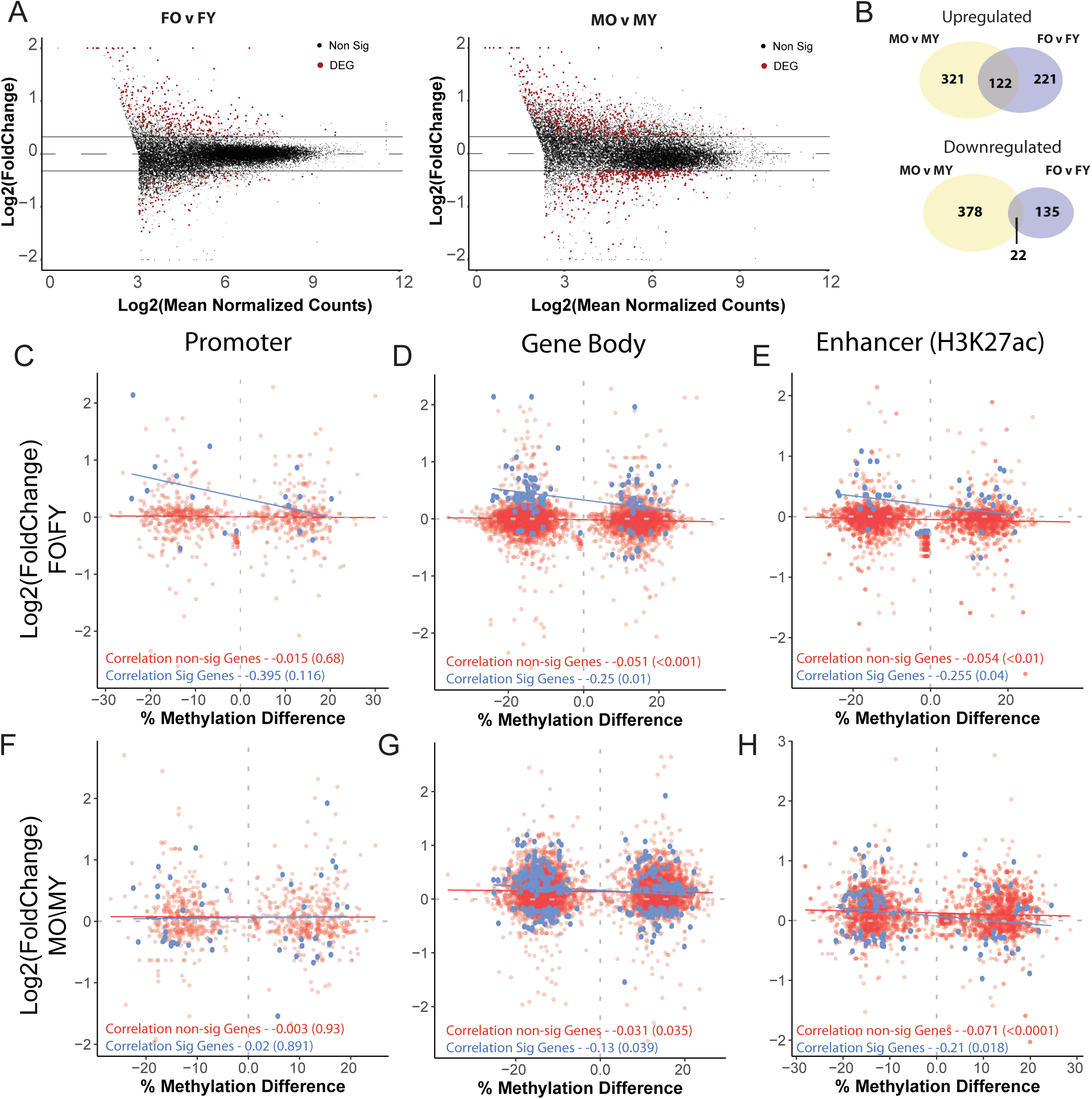
Differential methylation with aging is weakly anti-correlated with expression changes in gene body and enhancer regions. A) Volcano plots showing mRNA differential expression with aging (multiple linear regression, FDR < 0.05, |FC| > 1.25, n = 6/group) in males and females. B) Venn diagram showing the overlap of upregulated and downregulated differentially expressed genes between males and females. C-H) Correlation between age-DMRs mapped to promoters (C,F), gene body (D,G) or enhancer regions (E,H) and gene expression fold change (O/Y) in statistically significant (blue) and non-statistically significant genes (red) in females (C,D,E) and males (F,G,H).

To determine regions of differential methylation, the genome was binned to 500 bp non-overlapping windows. Windows with ≥ 10 CpGs and at least 3x coverage per CpG were retained yielding 979,603 regions analyzed for differential methylation with aging. Both males and females had roughly similar numbers of age-related differentially methylated regions (age-DMRs: 7702 in females vs 7029 in males) and showed a slight bias towards hypomethylation (Figure 1A,B). Only 2% of all age-DMRs were common to both males and females (Figure 1B). Of these sex-common changes, 68% were commonly regulated, e.g. hypermethylated in both males and females (χ2 test of independence p-value = 1.3 × 10^−6^). These results demonstrate that genome-wide, age-related changes in DNA methylation are predominately sex-specific, in agreement with prior findings [27].

Functional enrichment of genes containing age-DMRs revealed that although age-DMRs in males and females occurred in different genomic locations, genes containing age-related differential methylation are enriched in pathways with functional similarities, for example, genes containing age-DMRs in females are enriched in inositol phosphate metabolism, while genes containing age-DMRs in males are enriched in phospholipid metabolism and phosphoinositol metabolism. (Figure 1C,D, supplemental table 1). Generally, pathways common to both males and females are involved in glucose and lipid metabolism, neuronal interactions, and cellular integrity. These results suggest that while sex-divergence occurs at the level of the genome, the pathways affected by aging may still be functionally similar.

Age-DMRs were assessed for their enrichment across genomic features and gene regulatory elements. Over-representation of age-DMRs was observed in CpG islands and shelves, and within gene bodies (Figure 1E). Generally, DMRs were not enriched in promoter regions, but when separated according to their overlap with CpG islands, significant enrichment of age-DMRs is observed in promoters not associated with a CpG island. This is consistent with previous studies indicating that enrichment of DNA methylation occurs predominately outside of CpG islands associated with promoters [55,56]. Age-DMRs were also over-represented in active and poised distal gene regulatory regions, namely active enhancers and promoter flanks. This was also evident by over-representation of age-DMRs in hippocampal H3K27ac and H3K4me1, both indicators of active and poised enhancers [57,58] (Figure 1E). Hypomethylated age-DMRs were also over-represented in H3K36me3, a marker of exons and transcriptional elongation [59,60] shown to be altered with aging and associated with longevity [61,62], and in H3K27me3, a marker associated with gene repression (Figure 1E). Overall, enrichment of age-DMRs in genomic regions suggest that methylation of certain genomic regions is more susceptible to change with age as compared with others.

### Association between differential gene expression and differential methylation with aging

DNA methylation functions to modulate genomic architecture and regulate gene expression. However, the relationship of differential methylation to altered steady state gene expression with aging has not been comprehensively addressed. We used RNA-sequencing to analyze transcriptional differences with aging in the same samples used for methylation analysis to correlate age-DMRs with age-related differentially expressed genes (age-DEGs) in the hippocampus. With aging 781 genes were differentially expressed with aging in males and 433 in females (multiple linear regressions, fdr < 0.05 and |FC| > 1.25) (Figure 2A,B). Approximately 1/3 of the genes upregulated with aging were common between males and females (Figure 2B), and only 22 downregulated genes were common between the sexes (χ^2^ test of independence p-value < 2.2 × 10^−16^). This is consistent with previous findings reporting sexual divergence in transcriptional profiles in addition to a common core set of genes with aging[54].

In both males and females, only a small number of age-DEGs contained an age-DMR in their promoter region (±1kb of the TSS). The association between age-DMRs and differentially expressed genes with aging in promoters was not significant in both males and females (Figure 2C,F). When assessing all age-DMRs independent of their location in the gene body (TSS to TES), a weak negative correlation is observed in both males (r = −0.13, p = 0.039) and females (r = −0.25, p = 0.01) (Figure 2D,G). Given that DNA methylation can regulate gene transcription through changes in enhancer regions we examined the correlation between age-DMRs mapped to enhancer regions (determined by H3K27ac ChIP data from cortex) and transcriptional changes of their nearby genes. A significant negative correlation was observed between age-DMRs in enhancer regions and age-DEGs in both males (r = −0.21, p = 0.018) and females (r = −0.25, p = 0.04) (Figure 2E,H). Age-DMRs mapped to gene bodies or enhancers associated with genes that were not differentially expressed with aging resulted in significant, but very weak negative correlation (r < 0.1) in both males and females (Figure D,E,G,H). Taken together, age-DMRs may explain a small portion of the transcriptional changes that occur with age, and generally this effect is observed in enhancers and gene body, but not promoters. These findings are in agreement with recent studies in the liver showing a limited inverse association between gene body methylation with aging and gene repression of genes involved in lipid metabolism and growth hormone signaling [35]. Additionally, DNA methylation changes do not correspond with transcriptional changes in the CNS during neuronal maturation [43] or following induction of methylation in culture [63]. Therefore, while the canonical regulation of gene transcription by DNA methylation is likely to explain a small portion of the age-associated differential gene expression leaving age-related differential methylation to potentially serve a more complex role in transcriptional regulation than simply induction and suppression of steady-state gene expression.

### Age-related gene expression Changes are associated with young methylation profile

DNA methylation can play multiple roles in regulating gene transcription by altering protein binding occupancy [64], regulation of alternative splicing [65-69], and through interactions with histone marks [70,11]. To examine DNA methylation patterns between genes differentially expressed during aging and those that remain unaltered, correlations between fold mRNA change with aging and gene body methylation levels (mean methylation from TSS to TES) (Figure 3A,B) in early and late life were performed. Intriguingly, genes differentially expressed with aging show a moderate positive association between age-related mRNA expression change and gene body methylation levels, at both young and old age demonstrated by the regression lines in Figure 3A and 3B.Genes whose expression does not change with aging do not show an association (Figure 3A,B). That is, the average methylation levels of genes that were downregulated with aging was lower early in life, and remained lower throughout life as compared to genes that were upregulated with aging (Figure C,D).. This relationship was consistent in both young and old animals and was not influenced by age-related changes in methylation (Figure 3C,D). Although transcriptional changes with aging are predominately sex-specific, this association was evident in both males and females (Figure 3), with males showing a stronger association as compared to females.

**Figure 3:**
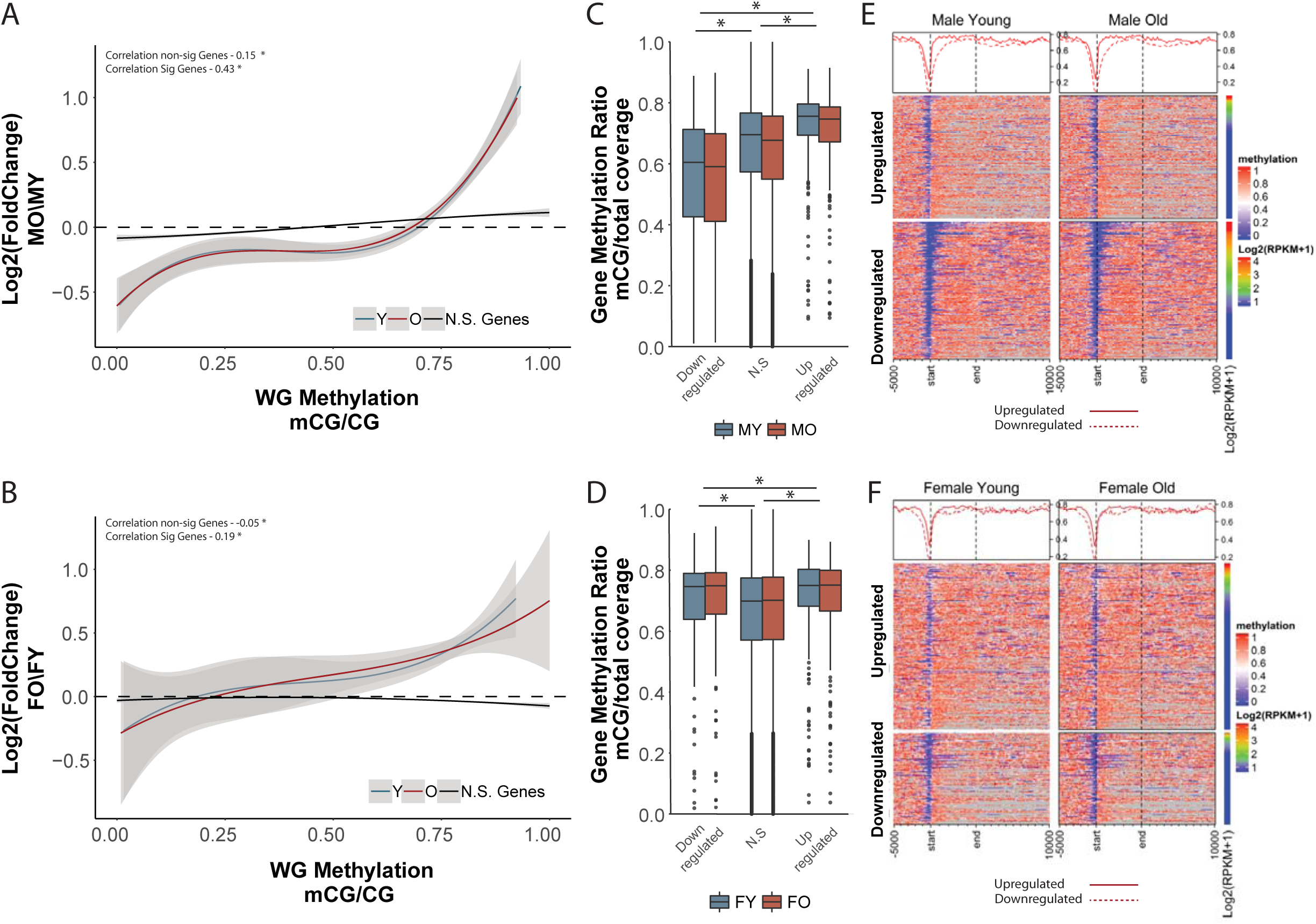
Direction of change of age-related differentially expressed genes is positively associated with gene body methylation. A,B) Genes downregulated with aging have lower gene body methylation at young age (blue regression line) in both males (A) and females (B) compared to genes upregulated with aging. This relationship is maintained with aging (red regression line). Curve corresponds to polynomial regression curve across significant (red and blue) and non-significant (black) differentially expressed genes, 95% confidence intervals are shaded by the grey area. Gene body methylation was calculated as methylation of all cytosines between the transcription start site and transcription end site of a given gene. C,D) Box plot of whole gene methylation grouped by genes upregulated, non-differentially expressed, and downregulated genes in males (C) and females (D). * p< 0.001 (Kruskal-Wallis Test) E,F) Heatmaps illustrating the per-gene gene body methylation patterns of genes upregulated and down regulated with aging in young and old, male (E) and female (F) animals.

Qualitative assessment of the DNA methylation landscape of up- and down-regulated genes with aging revealed that the main difference between up- and down-regulated genes occurs primarily around the transcription start site (Figure 3E,F). Therefore we repeated the analysis focusing on promoter methylation defined as +/-1kb of the TSS. The positive association between differentially expressed genes and baseline DNA methylation was recapitulated when examining only the promoter region (Figure 4A,B), and was comparable between sexes (Figure 4 C-F). Genes that did alter with aging showed a weaker association that was not consistent between males and females (Figure 4A,B).. The correlation between promoter methylation and gene expression changes was greater compared to that of expression with gene body methylation and was also independent of apparent age changes in methylation. Our observation reveals a relationship between age-related gene expression changes and DNA methylation that depends on the methylation patterns established early in life rather than differential methylation with aging. To determine whether the positive association between DNA methylation patterns and transcriptional changes with aging is observed in other tissues, we performed our analysis using paired WGBS and RNA-sequencing in the liver [35] (data obtained from GEO:GSE92486). A positive relationship between fold change and gene body methylation was observed with the liver data similarly to that observed in the hippocampus (Supplemental Figure 3). The lack of whole-genome bisulfite sequencing data with aging prevents the extension and validation of this analysis to other tissues.

**Figure 4:**
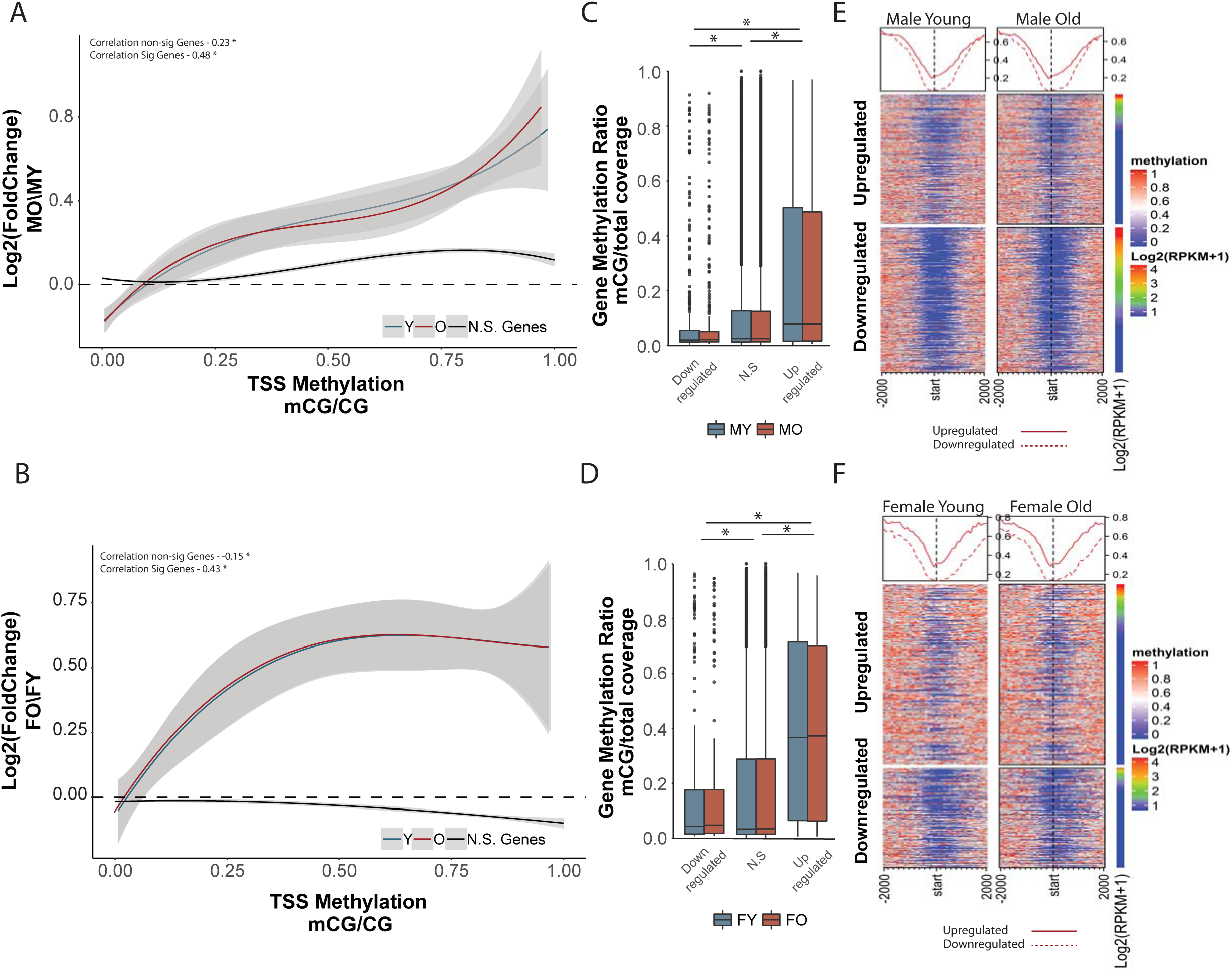
Direction of change of age-related differentially expressed genes is positively associated with promoter methylation. A,B) Genes downregulated with aging have lower promoter methylation at young age (blue) in both males (A) and females (B) compared to genes upregulated with aging. This relationship is maintained with aging (red). Curve corresponds to polynomial regression curve across significant (red and blue) and non-significant (black) differentially expressed genes, 95% confidence intervals are shaded by the grey area. Promoter is defined as ±1kb from transcription start site. C,D) Box plot of promoter methylation grouped by genes upregulated, non-differentially expressed, and downregulated genes in males (C) and females (D). * p< 0.001 (Kruskal-Wallis Test) E,F) Heatmaps illustrating promoter methylation patterns of genes upregulated and down regulated with aging in young and old in male (E) and female (F) animals.

### Association of methylation patterns with transcriptional changes with aging is not random

Differentially expressed genes with aging appear to have a different DNA methylation profile compared to genes that are stably expressed across the lifespan (Figures 3 and 4). To determine whether this observation is unique to genes that are differentially regulated with aging, we used a random sampling approach to correlate gene body DNA methylation values with their corresponding mRNA fold change with aging. Randomly sampled sets of 500 genes (n = 10,000) showed weak correlation (r < 0.1) compared to that observed for genes differentially expressed with aging (r > 0.4). This correlation was within the range of genes that were not differentially expressed with aging (Figure 5A).

**Figure 5:**
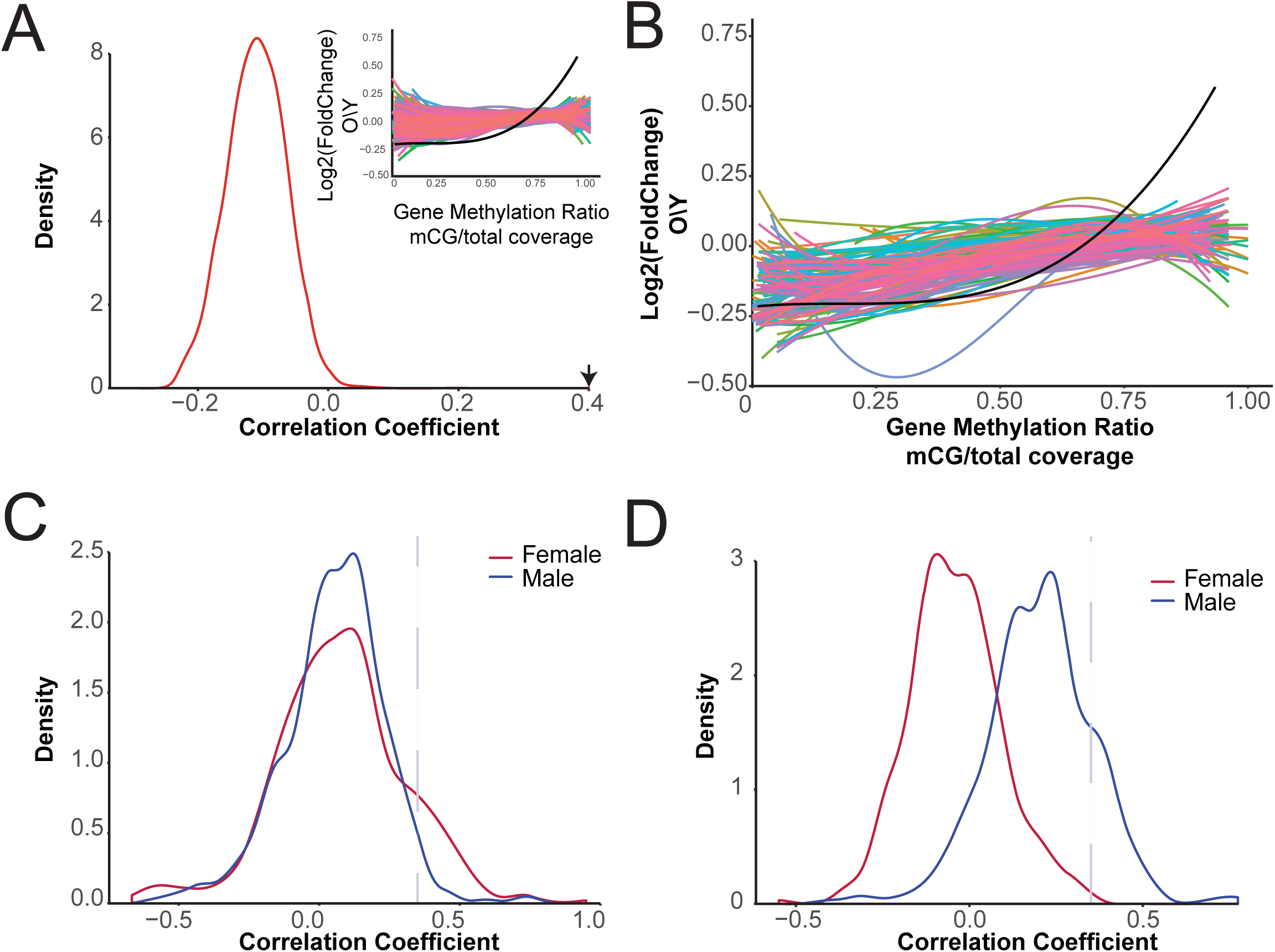
The association between differential expression and DNA methylation patterns in young animals is not random. A) Distribution of the correlation coefficients generated by correlating log2 fold change with gene body methylation of 500 randomly sampled genes (N = 10,000). Arrow indicates the location of the correlation coefficient of gene body methylation and differentially expressed genes in males. Snippet showing the polynomial regression curves of randomly selected gene sets compared to that observed in males (black regression line). B) Correlation between age-related differential genes expression and gene body methylation of Reactome pathways gene sets (only pathways with > 50 genes are included). Regression curve through all differentially expressed genes with aging and gene body methylation in males is shown in black. C,D) Distributions of the correlation coefficients generated by correlating log2 fold change with promoter (C) or gene body methylation (D) for each Reactome pathway gene set.

Next we asked whether genes sets that belong to the same pathway present a similar positive association. Pathways were extracted from the Reactome pathway database [71], and used as gene sets for correlation between methylation levels at young age and mRNA fold change with aging. After filtering pathways containing ≥50 genes, 368 pathways remained for the analysis (Figure 5B). Out of all the pathways analyzed, 35 pathways showed a correlation coefficient that met or exceeded the correlation coefficient of r > 0.4 (Figure 5C) observed between promoter methylation and genes differentially expressed with aging. For gene body methylation 32 pathways met or exceeded the correlation coefficient cutoff (Figure 5D) and were observed only in males. Pathways that showed the highest correlation between DNA methylation patterns and transcriptional change with age were pathways previously shown to be involved with aging included inflammatory pathways (transcriptional regulation by RUNX1, MHC II signaling, interferon signaling), oxidative stress, proteolysis, cell senescence, epigenetic regulation, and estrogen signaling (Supplemental Tables 2 and 3).

A central geroscience concept is that age-related changes intersect with those involved with disease pathogenesis, including Alzheimer’s disease [72,18]. Therefore we hypothesized that a positive correlation between transcriptional changes with neurodegeneration and DNA methylation profiles would be observed. In order to identify genes altered following neurodegeneration in the hippocampus, we used published RNA-sequencing data from two models of AD (APP and Ck-p25) and examined whether gene body and promoter DNA methylation levels in young and old animals are associated with differential gene expression observed in a neurodegenerative disease model. A significant number of genes were unique to each of the models; however significant overlap was observed between both AD models and with genes altered with aging (APP:Aging χ^2^ p < 2.2×10^−16^; CK-p25:Aging χ^2^ p < 2.0×10^−14^; APP:CK-p25 χ^2^ p < 2.2×10^−16^) (Figure 6A). As observed with genes differentially regulated with aging, upregulated genes with both APP and CK-p25 had significantly higher mean methylation in early life compared to downregulated genes (Figure 6B,C). Similar to differentially expressed genes with aging, this was observed for gene body (Figure D,F) and promoter methylation (Figure E,G). Differences in methylation in these models were not examined, therefore a potential difference in methylation due to AD pathology as a driving mechanism of differential gene regulation cannot be excluded, however, our findings suggest that genes differentially regulated following neurodegeneration may be more susceptible to change due to their methylation profile in a manner similar to that observed for genes differentially expressed with aging.

**Figure 6:**
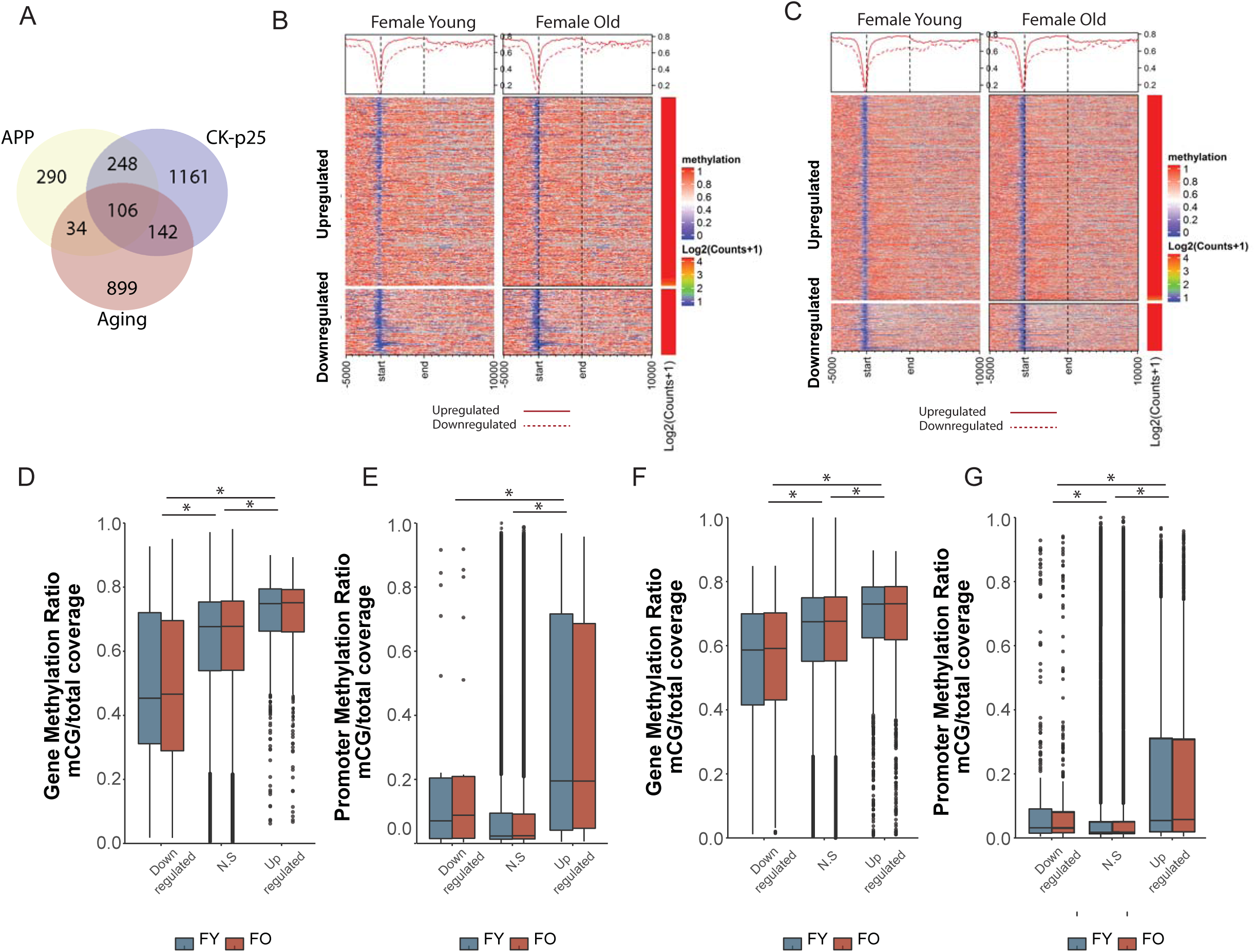
DNA methylation patterns in hippocampus of young and old animals is associated with genes differentially regulated in models of neurodegeneration. A) Venn-diagram representing the overlap between genes differentially expressed in two models of neurodegeneration (APP and CK-p25) and genes differentially regulated with aging (males and females combined). B,C) Heatmaps illustrating the per-gene gene body methylation patterns of young and old animals (females only) in genes upregulated and downregulated in two models of neurodegeneration (B. APP, C. CK-p25). D-G) Box plots of gene body (D,F) and promoter (E,G) methylation grouped by genes upregulated, unchanged or downregulated in APP (D,E) and CK-p25 (F,G).

### DNA methylation based prediction of differential expression with aging

To quantify the relative contribution of early life epigenetic patterns to predict gene expression changes with aging we used random forest (RF) modeling. The RF models were trained to predict the direction of transcriptional change with age (upregulated or downregulated) based on epigenetic marks annotated in the hippocampus and cortex obtained from publicly available datasets (see methods). The epigenetic marks used to predict differential expression included histone breadth of coverage (breadth/gene size) of H2Bac, H3K27ac, H3K27me3, H3K36me3, H3K4me3, H3K9me3, H2A.Z (both young and old), gene body DNA methylation in young and old, promoter methylation in young and old, gene size, and baseline expression defined as mean expression levels or RPKM of a gene in young animals.

The trained RF model was able to correctly classify transcriptional changes with high accuracy in both males (83%) and females (77%) (Figure 7A,B). RF performance decreased slightly when trained based on DNA methylation means and RPKM alone, but still performed significantly better than random in both males (78%) and females (70%) (Supplemental Figure 4). Evaluation of feature importance to each of the RF models revealed that DNA methylation and gene size are highly important for predicting gene expression in both males and females. In males, gene size, H2A.Z marks, H3K4me3, H3K27ac, and DNA methylation averages of both whole gene and promoters (Figure 7C) contributed the most to predictive accuracy. In females, high importance features for model prediction included mean expression, DNA methylation levels and gene size (Figure 7D). While H3K27ac, H2A.Z and H3K4me3 were still important for model prediction, their weight in predication accuracy decreased compared to that observed in males. The differences in feature importance between males and females may be a result of sex-differences in epigenetic landscape that is not captured by the GEO datasets used in the current analysis which were generated from males.

**Figure 7:**
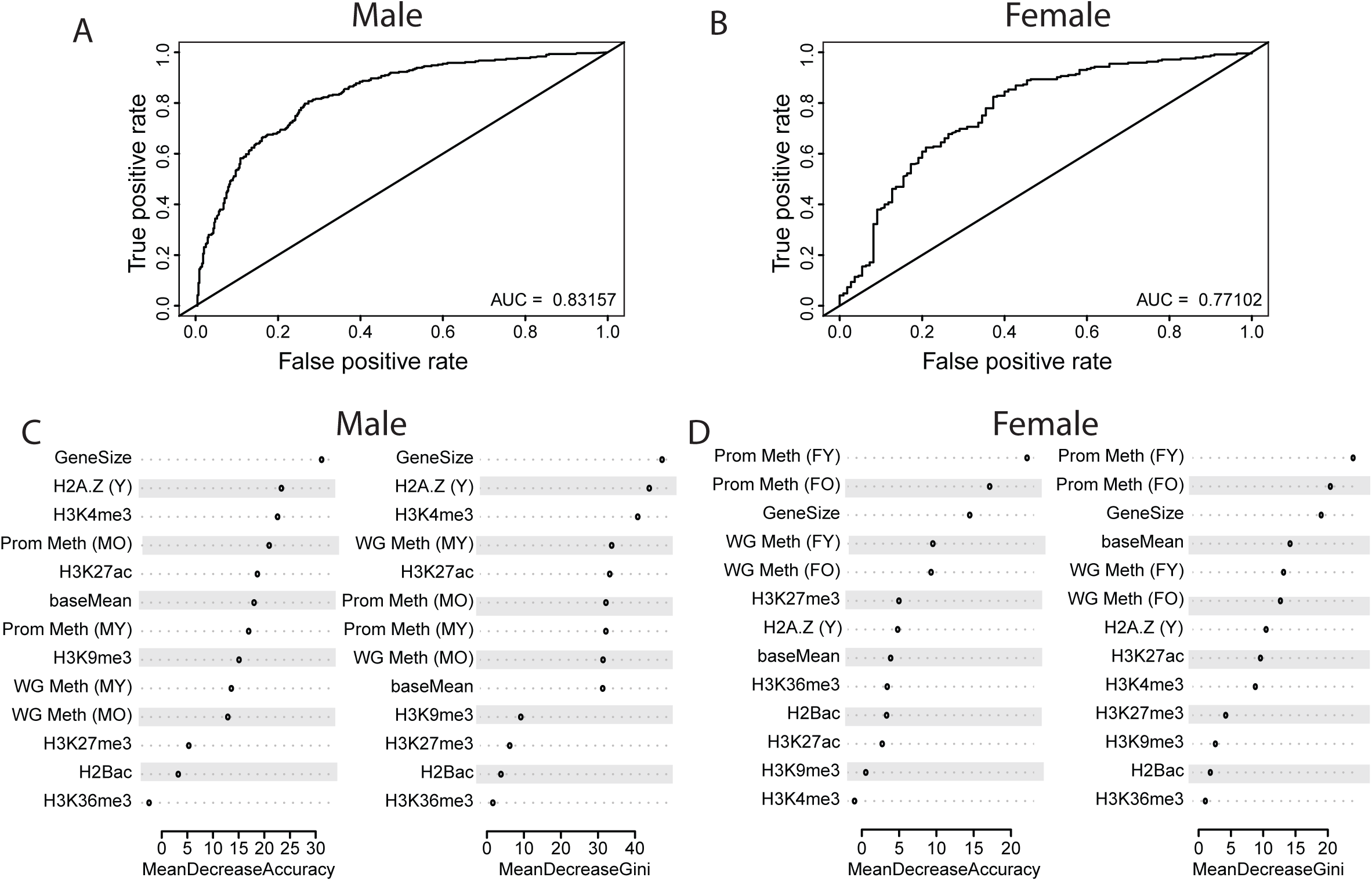
Direction of change of age-related differentially expressed genes can be predicated based on epigenetic marks in young age. A,B) Area under the curve of the receive operating characteristic (ROC) curve showing the classification accuracy of differentially expressed to upregulated and downregulated genes for Random Forest model in males (A) and females (B). C-F) Feature importance of epigenetic marks for classification accuracy (mean decrease accuracy and mean decrease gini) in males (C,D) and females (E,F).

It should be noted that these different epigenetic marks are not independent of each other as DNA methylation is closely associated with both H3K4me3, an active promoter mark [73], and H3K27ac, an enhancer mark [74]. Regions of H3K4me3 and H3K27ac often act coordinately with DNA methylation during gene transcription regulation [75]. Local depletion of DNA methylation is a hallmark of H3K4me3 and H3K27ac [58], and thus these marks are considered to be regulated by DNA methylation. Gene size was a significant contributor to the accuracy of the models (Figure 7C,D) and is also regulated by DNA methylation. For example, in the CNS epigenetic regulation of long genes by DNA methylation often occurs developmentally through regulation of the DNA methylation binding protein MeCP2 [76]. Our findings are in agreement with those of Benayoun et al. [77] which examined some of these marks but not DNA methylation in the cerebellum and olfactory bulb. Taken together, our result put forward the idea that epigenetic regulation at a young age may direct transcriptional change with aging.

## Discussion

By analyzing the methylation and transcriptional profiles in the hippocampus of young and old animals we provide evidence for a potentially novel role for DNA methylation in regulating transcriptional changes with age that is independent of age-related changes to the methylome. Additionally, differences in methylation with age are enriched in exonic and intronic regions, and showed a weak inverse correlation with differences in gene expression. The functional role of gene body methylation is not yet well defined but is associated with transcriptional elongation [78], splicing [66,67,79,69], regulation of alternative promoters [80], and modulation of expression levels through interaction with methyl binding proteins such as MeCP2 [81,82]. In the CNS, in contrast to other tissues, gene body methylation is inversely correlated with expression levels [1,83]. Our findings are consistent with this finding. However, the diverse functional roles of gene body methylation create a challenge in interpretation of the association between gene body age-DMRs and the altered transcriptional profile with aging. Nevertheless, age-related differential methylation within genes is common to various tissues; therefore improved knowledge on how gene body methylation regulates expression is required in order to understand the potential functions age-DMRs play in regulation of the aging transcriptome.

### Association of promoter age-DMRs with age-DEGs is limited

The association between DNA methylation and gene expression is often derived from the inverse correlation between mRNA expression and DNA methylation in promoters under normal conditions [8]. While differences in promoter methylation in the hippocampus occur with aging, the genes associated with these promoters are generally not differentially expressed with age (Figure 2). A potential explanation is that observed changes to the methylome with age are subtle and therefore insufficient to induce transcriptional differences, however a weak correlation between gene expression changes and differential promoter methylation is also observed in studies of cancer and cellular differentiation [84,85], which include disruption to- (cancer) or reprogramming of-(differentiation) the methylome. The limited correlation between age-related differential promoter methylation and gene expression changes does not preclude differential promoter methylation from altering expression of specific genes, but is insufficient to explain the majority of transcriptional changes observed with age in the hippocampus. It should also be noted that gene expression changes rapidly with stimuli and the expression levels here were collected to represent steady-state expression levels.

### Enhancer age-DMRs are related to age-DEGs

Recent studies identified that altered DNA methylation patterns play a greater role in explaining transcriptional changes when they occur in distal regulatory regions, namely enhancers, compared to gene promoters [85]. Age-related differential methylation is enriched in enhancer marks in various tissues [86,87,39,88], including in the hippocampus [26]. With aging, altered methylation in differentiating cells, specifically hypomethylation, was shown to be enriched in regions marked by H3K4me1 [89], a marker of active and poised enhancers [90], and is thought to activate gene expression. Consistent with these findings, we found enrichment of both hyper- and hypo-methylated age-DMRs in regions distal to gene promoters, specifically in annotated active and poised enhancers. These age-DMRs were inversely correlated with transcriptional differences with aging in both males and females.

Recent findings shed light on the interaction between the enhancer marks H3K27ac and H3K4me1 and DNA methylation and the functional role of this interaction on gene transcription regulation [91]. Enhancer activation can be both positively or negatively associated with DNA methylation depending on the regulatory nature of the enhancer and the developmental stage of the organism [57,58,92]. Enhancers containing transcription factor binding motifs tend to be inversely correlated with DNA methylation late in life, but not during cellular differentiation where DNA methylation increases in enhancers proximal to genes that involve cellular specification [75]. Methylation of super-enhancers is thought to contribute to the structural integrity of the genome at these regions [93,92]. Although alterations in chromatin landscape with aging have been reported, few studies have mapped altered histone marks with age. H2A.Z, a histone variant needed for the acetylation of histone 3 lysine 27 [94], do change with aging in the hippocampus [95], and may be a contributing mechanism to enhancer mark changes with aging. Given these results we hypothesize that changes in methylation can potentially alter transcription through attenuation of enhancer strength rather than facilitating deposition of H3K27ac. Future studies are required to address this hypothesis by mapping the differences in enhancer landscape with age in both male and female and in different cell/tissue types.

A unique feature of genes that were differentially expressed with age was their association with DNA methylation patterns established in early life (i.e. methylation levels in young animals). Levels of methylation of upregulated genes was higher than that of downregulated genes in young animals, this difference persists in old animals and therefore was generally independent of age-related differential methylation. The association between methylation levels and expression was not observed for genes that were not altered with aging or randomly selected genes. Furthermore, gene expression changes with aging were generally different between males and females, yet a similar association was observed in both sexes. This finding supports the concept that, based on their epigenetic patterns established early in life, specific genes have a higher propensity to change with age than others and that their induction/reduction is dependent on the methylation status of the gene. Therefore suppression or induction of genes with aging is likely to occur downstream of methylation by factors that interact with the methylome such as histone modifications or methyl binding protein dynamics. An additional finding was that genes that changed with age and correlated with early life methylation occur in specific gene sets that function in similar pathways. This is consistent with the notion that genes with similar functions are regulated in similar ways [96,97].

Using the predictive capabilities of machine learning we were able to show that baseline expression and DNA methylation levels alone can classify whether differentially expressed genes will be downregulated or upregulated. When other epigenetic marks from the young/adult brain are added to the model, the classification accuracy of the model improves. This provides further support to the idea of early epigenetic programing as a determining factor of expression changes with age. A recent study [77] showed similar results by predicting age-related expression changes based on chromatin marks. The authors found that changes in the enhancer mark H3K27ac with age were among the highly important features for accurate classification. This indicates that age-related alterations to the epigenome contribute to transcriptional changes with age. Although changes in chromatin predict gene expression changes well, we were able to achieve similar predictive capabilities based on early life DNA methylation alone. Future studies combining both baseline epigenetic profiles and age-related alteration to histones are needed to improve the classification accuracy of these models, and perhaps help identify the mechanisms that underlie epigenetic regulation of transcriptional changes with aging.

The processes underlying aging are thought to promote the development of age-related neurodegenerations [49]. In our study we find that the association between early life methylation patterns and differential gene expression is also observed in genes that are dysregulated in mouse-models of AD. That is, genes that were upregulated in a model of neurodegenerative disease had higher gene body methylation at young age compared to those that were downregulated. Thus it is plausible that DNA methylation patterns at young age may account for transcriptional changes that facilitate the development of more severe conditions through altered gene expression late in life. Additionally, variation in transcriptional profiles with aging was reported between mouse strains from diverse genetic backgrounds [98], signifying the importance of the genome in determining transcriptional regulation with age. Genetic factors are also thought to underlie increased longevity in supercentenarians [99,100] and may contribute to age-related epigenetic drift [101]. Therefore, population differences in methylation early in life, in addition to genetic diversity may be associated with the variability in development of age-related neuropathologies, predisposition, and severity observed between individuals.

Additionally, aging rate can also be influenced by environmental factors which influence epigenetic programming. For example early life adversities have been demonstrated to cause alteration to the epigenome, result in accelerated epigenetic aging, and can persist in late-life [102-104]. Therefore, epigenomic programming during developmental stages and early adulthood may serve as a potential mechanism for altered late-life outcomes, including aging and susceptibility to disease. In addition, DNA methylation patterns are also altered with anti-aging therapies that have a beneficial effect on molecular and cellular aging hallmarks [39,26]. These therapies, for example calorie restriction, have been shown to be potent in a short window early in life and following life-long treatment [105,106]. A point to further investigate is how these anti-aging therapies can alter methylation patterns both early and late in life to prevent age-related transcriptional changes and promote a pro-longevity phenotype.

### Conclusions

Age-related differences in epigenetic marks are likely to contribute to transcriptional alterations, however, these epigenetic differences account for a small subset of the gene expression changes with aging and are dependent on the genomic location, e.g. promoter vs. regulatory region. It is noteworthy that our current knowledge of the exact location of regulatory marks is far from complete and is likely to vary between cell types and tissues, therefore without the ability to map the exact location of a TSS, alternative splice sites, gene regulatory marks and the genes they interact with, findings should be interpreted with caution. Our current findings identify a potential new way in which DNA methylation can influence age-related transcriptional change. The early establishment of DNA methylation patterns of a gene appears to partially determine whether the gene will change with age and the directionality of the change. Interestingly, a recent preprint identified a similar finding examining histone modifications in the cerebellum and olfactory bulb [77]. This association is also found in the liver and in differential gene expression in Alzheimer’s models. Together, these findings indicate that the early-life epigenetic landscape of a gene may direct its gene expression trajectory with aging and age-related disease. These findings provide a potential mechanism for the developmental origins of disease concept [107].

## Materials and methods

### Animals and nucleic acid extraction

Male and female C57BL/6 mice were obtained from the NIA aging colony at 2 and 21 months of age. Mice were housed at the University of Oklahoma Health Sciences Center barrier animal facility and maintained under SPF conditions until 3 and 24 months of age. All experimental procedures were performed according to protocols approved by the OUHSC Institutional Animal Care and Use Committee. Mice were euthanized by decapitation and hippocampal tissue was dissected and snap frozen until used for DNA and RNA extraction. DNA and RNA from young and old animals (n=6/group) was isolated from hippocampal tissue using Zymo Duet DNA/RNA (Zymo research).

### Whole Genome Bisulfite Sequencing and DMR calling

Isolated genomic DNA from young and old animals (n=3/group) was used for Whole-Genome Bisulfite Sequencing (WGBS). Bisulfite conversion was carried out using EZ DNA methylation Lighting (Zymo Research, Irvine, CA) and library construction used Swift Accel-NGS methyl-seq kit reagents (Swift Bioscience, Ann Arbor, MI) following manufacturer’s instructions. Library size was assessed by Tapestation (Agilent Technologies, Santa Clara, CA) and quantified by quantitative PCR (Kappa Biosystems, Wilmington, MA) prior to sequencing. BS-seq libraries were sequenced by 100 bp paired-end reads on the Illumina HiSeq-2500 (Illumina, San Diego, California, USA).

Paired-end reads were trimmed using trimmomatic version 0.35 [108]. Reads were adapter trimmed and filtered based on quality. Bases with a Q-score < 30 were removed from the 5’ and 3’ ends. Reads were quality filtered using a sliding window approach (parameters were set to 5:30). Additionally, reads shorter < 25bp post-trimming were removed. Trimmed PE reads were aligned to the mouse reference genome (GRCm38/mm10) with Bismark Bisulfite Mapper version 0.14.4 [109] using default settings. Methylation % and coverage of each CpG site were extracted with bismark methylation extractor. Mean Coverage per sample was 5x (±0.4 SD), sites with < 3x mean coverage per group were removed resulting in >20 million CG sites analyzed.

In order to determine differentially methylated regions (age-DMRs), the genome was binned into consecutive, non-overlapping 500 bp windows. Samples within each group were combined to achieve higher coverage per site, and windows with <10 CpG sites were omitted from the analysis. Statistical significance of differential methylation was determined using Fisher exact test followed by false-discovery multiple testing correction. Differentially methylated regions were considered statistically different if FDR adjusted p-value <0.05.

### RNA sequencing and differential gene expression analysis

RNA integrity was quantified by TapeStation (Agilent Technologies, Frankfurt, Germany) and samples had RNA integrity numbers >8. RNA-sequencing libraries were prepared using Illumina’s TruSeq RNA-seq library prep with a rRNA depletion step according to manufacturer’s instructions. Libraries were sequenced with 150bp paired-end (PE) reads on the Illumina HiSeq 3500 platform (Illumina, San Diego, California, USA) (n =6/group). Sequence quality control was performed with fastQC. Following QC step PE reads were trimmed similarly to the WGBS sequences using trimmomatic.

Following QC and trimming, reads were aligned to the mouse (mm10) reference genome using STAR [110]. For alignment, the genome was prepared based on GENCODE M15 release. STAR Alignment parameters were set to: outFilterScoreMin 2, outFilterMultimapNmax 5, outFilterMismatchNmax 10, outFilterMatchNmin 20, outSJfilterReads Unique, outSJfilterOverhangMin 25 10 10 10, alignSJoverhangMin 2, alignSJDBoverhangMin 2, chimSegmentMin 25. Reads per gene were counted in R using the ‘summarizeOverlap’ function in the GenomicAlignments package. Raw reads were normalized using DESeq2 R package [111] and transformed using variance stabilized transformation. Differential expression between all groups was assessed using multiple linear regression (R package ‘glm’) using read counts as the dependent variable and age (young and old) and sex (male and female) as the independent variables. Genes with significant age main effect (p < 0.05) were then carried on for pairwise comparisons using Conover post hoc test followed by false discovery rate adjustment using ‘fdr’ as implemented in the R package ‘lsmeans’.

### Enrichment Analysis

For pathway enrichment age-DMRs were annotated using ChIPseeker [112], and enrichment analysis was performed using the R package ‘ReactomePA’ [113]. To determine over- and under-representation of DMRs in genomic features, annotated introns, exons, and CpG islands were obtained from UCSC Genome Browser. Promoters were defined as +/− 1kb from the transcription start site. CpG shores were defined as 2kb upstream and downstream of the annotated CpG island boarders and CpG shelves were defined as 2kb upstream and downstream from shores. Gene-regulatory regions in the mouse brain were extracted from Ensemble open database [114]. DMRs were mapped to genomic features using ‘bedtools’ [115]. Statistical significance of over- or under-representation of DMRs in genomic features was determined using hypergeometric test in R.

### Differential Expression Prediction

Differentially expressed mRNAs with aging were classified based on the directionality of change (upregulated or downregulated) and divided into a training set and a validation set by randomly subsetting 70% of the genes to the training set, the remaining genes were used for model validation. Prediction of gene change directionality with aging was performed separately for male and females. Random forest (RF) was used for prediction, and all analysis and cross-validation was performed in R using the ‘randomforest’ package. The RF model was trained based on selected epigenetic features including mean gene DNA methylation in young and old, mean promoter (+/-1kb of TSS) methylation in young and old, gene size, base expression at 3 months, and breadth of coverage of the following histone marks: H2A.Z from young and old animals, H3K27ac, H3K36me3, H3K4me3, H3K27me3, H2Bac, and H3K9me3. Breadth of coverage was calculated by the breadth sum of all peaks in a gene/gene length.

### Public Data Acquisition

Paired methylation and differential expression data for liver was obtained from GEO:GSE92486 [35]. Differential genes expression for age-related neurodegenerative disease APPswe/PS1δE9 (APP) and Ck-P25 models were obtained from GEO:GSE93678 [116] and GEO:GSE65159 [117]. Only WT control and experimental groups were used. ChIP-sequencing data of hippocampal histone marks was obtained from GEO:GSE85873 (H3K4me3 and H3K27me3) [118], GEO:GSE103358 (H2Bac), and GEO:GSE100039 (H2A.Z)[95]. Cortex epigenetic marks including H3K27ac, H3K36me3, and H3K9me3 were obtained from GEO: GSE103214 [119]. Peak calling was determined with MACS2 [120].

## Acknowledgments

The authors would like to acknowledge the University of Oklahoma Supercomputing Center for Education & Research (OSCER) for allocating computational resources used for data analysis and the Oklahoma Medical Research Foundation for sequencing assistance. This work was supported by Veterans Administration (I01BX003906), National Institute on Aging (P30AG050911, R56AG059430, T32AG052363), National Institute of General Medical Sciences (P20GM125528) and Oklahoma Center for Adult Stem Cell Research funding.

NH performed all post alignment data analysis. DRM performed the library preparations and file processing was performed by NH and DRS. Annotation of transposable elements was performed by LBB. WMF supervised all the work and provided funding. NH and WMF wrote the manuscript. All authors contributed to the interpretation of results and to the review and editing of the manuscript.

---

**Supplemental Figure 1.**
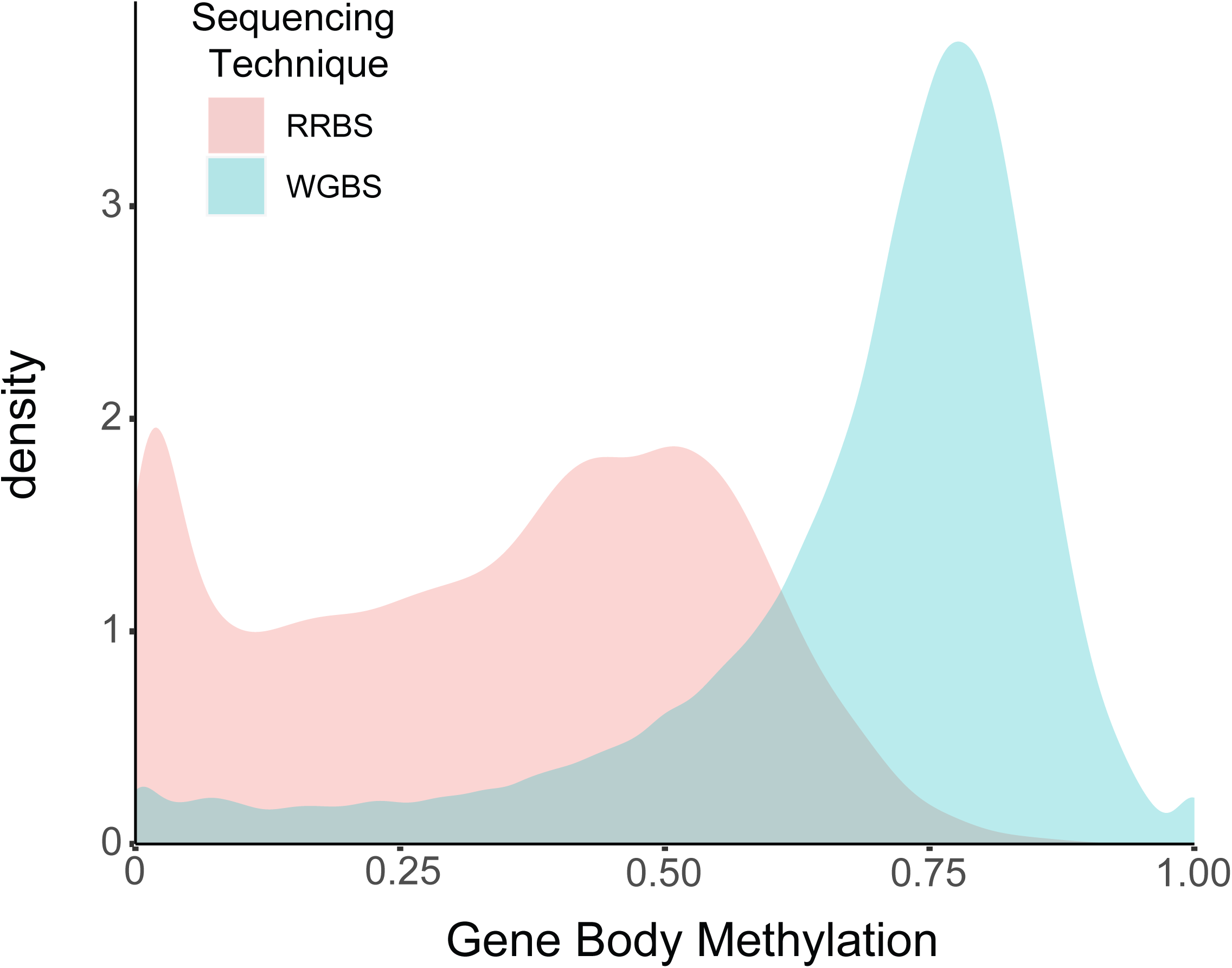
A comparison of the distribution of gene body methylation across all genes in the liver measured by reduced representation bisulfite sequencing (RRBS) and whole genome bisulfite sequencing (WGBS) obtained from GEO:GSE92486. RRBS covers methylation over 23000 genes as compared to 29000 by WGBS. The gene body methylation profiles obtained by RRBS do not represent the gene body methylation values observed by WGBS.

**Supplemental Figure 2.**
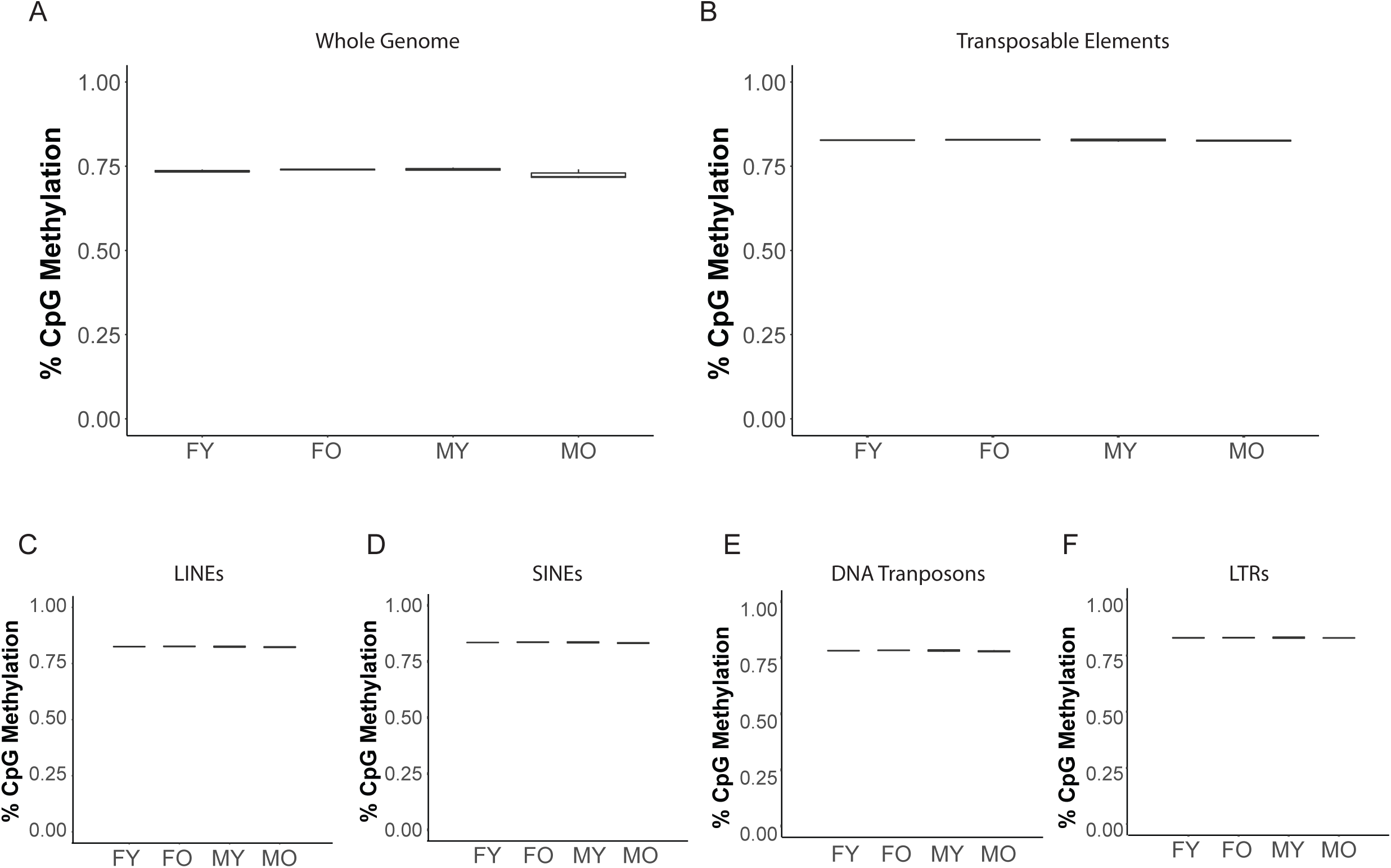
A) Box plots of whole genome methylation in young and old males and females. B-F) Box plots showing the methylation levels of cytosines mapped to all transposable element regions (B) or to specific transposable element families. Long interspersed nuclear repeats, LINEs (C), small interspersed nuclear repeats, SINEs (D), DNA transposons (E), and long terminal repeats, LTRs (F), n=3/group.

**Supplemental Figure 3.**
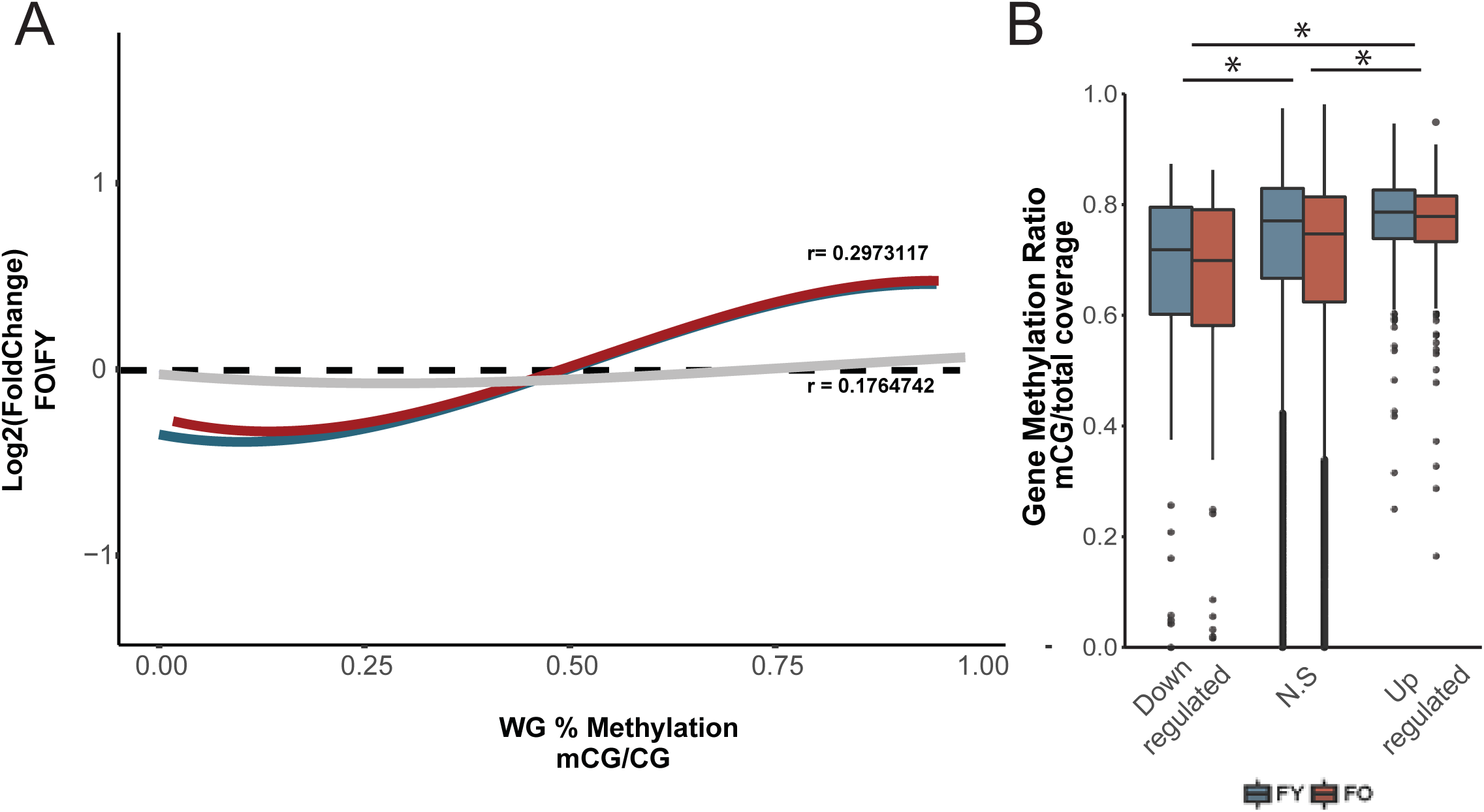
A) Genes downregulated with aging have lower gene body methylation at young age (blue regression line) compared to genes upregulated with aging in the liver. This relationship is maintained with aging (red regression line). Curve corresponds to the polynomial regression curve across significant (red and blue) and non-significant (black) differentially expressed genes. B) Box plot of whole gene methylation grouped by genes upregulated, non-differentially expressed, and downregulated genes in the liver. * p< 0.001 (Kruskal-Wallis Test).

**Supplemental Figure 4.**
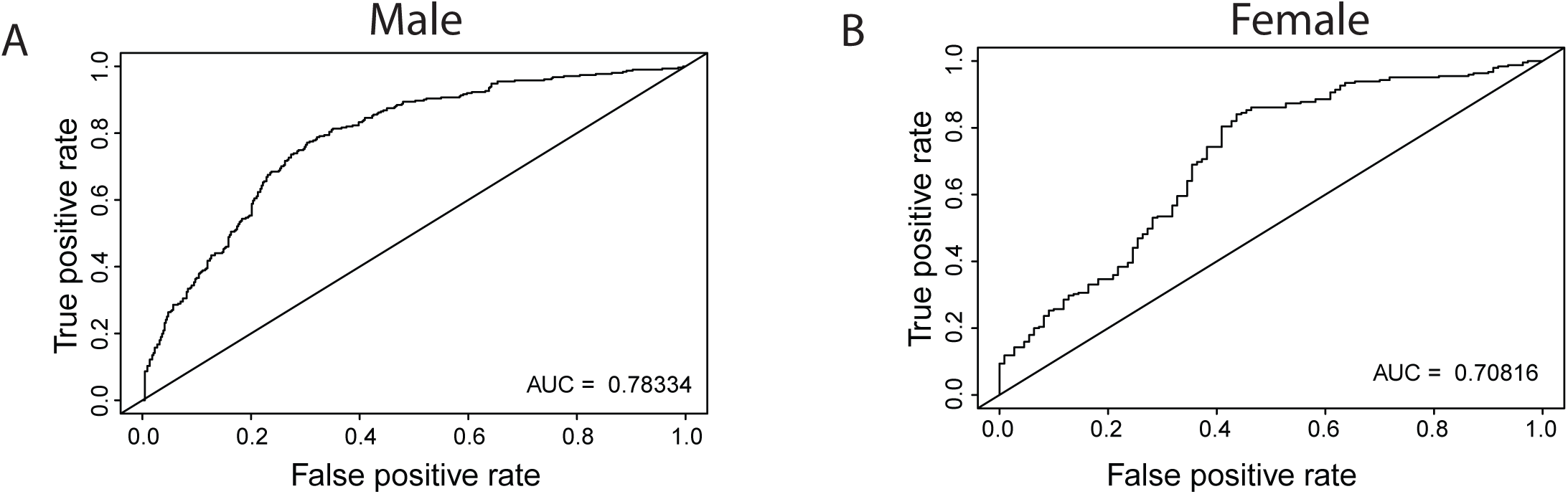
A,B) Area under the curve of the receive operating characteristic (ROC) curve showing the classification accuracy of age-related differentially expressed genes to upregulated and downregulated genes for Random Forest model in males (A) and females (B) trained based on baseline methylation and promoter and gene body DNA methylation.

